# Rab10 and Arhgef7 interact to promote axon extension by linking vesicle supply and exocytosis

**DOI:** 10.64898/2026.11.04.764424

**Authors:** Federica Olocco, Elena Bekker, Priyadarshini Ravindran, Thorben Pietralla, Andreas W Püschel

## Abstract

**Summary blurb:** The interaction of Rab10 and Arhgef7 is shown to promote axon extension by linking vesicle supply and exocytosis. Rab10 activates the Cdc42 GEF Arhgef7 on vesicles, which promotes membrane expansion.

The formation of functional neural circuits begins with the extension of a single axon by immature neurons. Axon elongation requires an expansion of the plasma membrane that depends on the polarization of vesicle trafficking and exocytosis. It is not completely understood how the increased transport of vesicles into the growing axon is linked to their subsequent insertion into the plasma membrane. An analysis of early stages of neuronal differentiation indicates that an asymmetry in neurite length precedes axon specification and the accumulation of PPVs in the nascent axon. We show that increasing the transport of vesicles in unpolarized neurons is sufficient to promote the extension of additional axons. This effect indicates that an increased supply of vesicles stimulates their subsequent insertion into the membrane to allow neurite growth. The link between vesicle transport and exocytosis is mediated by the interaction of Rab10 with Arhgef7. Rab10 stimulates Arhgef7 activity on PPVs as a GEF for Cdc42b to promote axon extension through exocyst-dependent exocytosis, suggesting that their interaction links the increased supply of PPVs to their exocytosis.

## Introduction

The highly polarized morphology of neurons with molecularly and functionally distinct axons and dendrites enables the formation of complex neuronal networks. Before immature neurons become polarized by extending a single axon, they first form several neurites that exhibit cycles of elongation and retraction (Dotti *et al*, 1988; Lin *et al*, 2026). Initially, all neurites have the potential to become an axon. This transient, unpolarized state is maintained by a balance of positive signals that promote neurite growth and negative signals that inhibit it (Namba *et al*, 2015; Schelski & Bradke, 2017; Yogev & Shen, 2017; Takano *et al*, 2019). An oscillatory process initiates actin waves that travel anterogradely along the neurite to the growth cone and promote neurite extension by opposing actin-dependent inhibitory signals (Flynn *et al*, 2009; Winans *et al*, 2016; Woo *et al*, 2019; Lin *et al*, 2026). This mechanism ensures that only a single neurite extends at a time (Lin *et al*, 2026). Eventually, one of the minor neurites begins to elongate continuously as the axon while the remaining minor neurites maintain their size until they differentiate into dendrites at a later stage (Dotti *et al*, 1988).

Pathways that modulate the dynamics of microtubules and actin filaments are critical regulators of axon extension (Burridge & Wennerberg, 2004; Alberts *et al*, 2006; Witte *et al*, 2008; Katsuno *et al*, 2015; Schelski & Bradke, 2017; Woo *et al*, 2019; Li *et al*, 2021; Burute *et al*, 2022; Schelski & Bradke, 2022; Lin *et al*, 2026). In addition, intracellular transport and exocytosis are essential to promote extension by providing material for the expansion of the plasma membrane (Lockerbie *et al*, 1991; Futerman & Banker, 1996; Bradke & Dotti, 1997; Tao *et al*, 2019). Membrane expansion is mediated by plasmalemmal precursor vesicles (PPVs), a specific type of post-Golgi vesicles in neurons (Lockerbie *et al*, 1991; Pfenninger *et al*, 2003; Pfenninger, 2009). PPVs are characterized by the presence of the small GTPase Rab10 and the SNARE protein VAMP4 (Dupraz *et al*, 2009; Pfenninger, 2009; Wang *et al*, 2011; Grassi *et al*, 2015). Rab10 links PPVs to Kinesin-1 that transports them into the axon, where they fuse with the plasma membrane of the growth cone (Wang *et al*, 2011; Deng *et al*, 2014). In the growth cone, PPVs are tethered at the plasma membrane by the octameric exocyst complex prior to their exocytosis, which is essential for membrane expansion (Craig *et al*, 1995; Lalli, 2009; Fujita *et al*, 2013; Deng *et al*, 2014; Wu & Guo, 2015; Bustos Plonka *et al*, 2022). Tethering and exocytosis of PPVs are regulated by several GTPases including TC10 and the brain-specific Cdc42 isoform Cdc42b that interact with the exocyst complex (Inoue *et al*, 2003; Dupraz *et al*, 2009; Ravindran & Püschel, 2023). Both GTPases are activated by the GEF Arhgef7, which is required for axon formation in cultured neurons and in vivo (López Tobón *et al*, 2018). The loss of Rab10, Arhgef7, Cdc42b or Exo70 blocks PPV exocytosis and axon formation (Wang *et al*, 2011; López Tobón *et al*, 2018; Ravindran & Püschel, 2023).

Vesicles are transported bidirectionally in axons and dendrites (Lasiecka & Winckler, 2011). Their antero- and retrograde transport remains balanced in unpolarized neurons until a nascent axon emerges (Bradke & Dotti, 1997; Pfenninger, 2009; Humpert *et al*, 2021). With the extension of the axon, vesicle transport becomes polarized and the exocytosis of PPVs in the axonal growth cone promotes axon elongation (Pfenninger & Friedman, 1993; Bradke & Dotti, 1997; Schelski & Bradke, 2017). Simulations based on a model for vesicle transport in polarizing neurons suggest that an increased exocytosis of vesicles together with the bidirectional transport between growth cones and soma can itself already result in a preferential trafficking of vesicles to the longest neurite (Humpert *et al*, 2021). This preferential transport into the longest neurite happens once a sufficient difference in neurite length is established. A biased distribution of vesicles to the longest neurite would, thereby, reinforce an initial asymmetry in neurite length and ensure the continued growth of the longest neurite while minor neurites remain unable to extend. An increased supply of PPVs to the longest neurite can promote extension only when it is linked to an enhanced exocytosis, leading to an increase in membrane insertion. However, a molecular mechanism that mediates a link between the transport of PPVs and their exocytosis remained to be identified.

Here, we show that an asymmetry in neurite length precedes axon specification, which coincides with an accumulation of PPVs in the nascent axon. Increasing the transport of PPVs into neurites is sufficient to promote the extension of additional axons. We identify an interaction of Rab10 and Arhgef7 that activates Cdc42b. The activation of Rab10 or Cdc42b by their respective GEFs specifically on PPVs promotes the extension of supernumerary axons while activation at the plasma membrane does not. Arhgef7 acts upstream of Cdc42b and the exocyst subunit Exo70 to promote axon extension. These results suggest that the interaction of Rab10 and Arhgef7 on PPVs promotes exocytosis by activating Cdc42b and, thereby, links vesicle transport to exocyst-dependent membrane insertion, which enables axon elongation.

## Results

### PPVs accumulate in the longest neurite

The continued growth of a neurite as it extends as an axon requires the expansion of the plasma membrane and depends on an increased supply of vesicles (Futerman & Banker, 1996; Pfenninger, 2009). To determine how the distribution of PPVs changes during the establishment of neuronal polarity we analyzed neurons at 12 h, 24 h, 36 h, 48 h, 60 h and 72 h of culture by staining with anti-acetylated tubulin and anti-VAMP4 antibodies as markers for axons and PPVs, respectively (Fig 1A). An asymmetry between neurites was first detectable at 36 h when one of the neurites became significantly longer. The difference in length between the longest and second longest neurite increased from 6.8 ± 0.9 µm at 12 h to 16.6 ± 2.6 µm (24 h) and 38.7 ± 3.8 µm at 36 h (Fig 1B). To determine at which stage VAMP4 begins to show a polarized distribution we calculated the polarity index as the ratio of the fluorescence intensity in the longest and second longest neurite. The restriction of acetylated tubulin to a single neurite was first observed between 36 h and 48 h (polarity index 36 h: 0.9 ± 0.1, 48 h: 1.7 ± 0.1; Fig 1C). At the same time, a significant increase in the signal for VAMP4 was observed in the axon compared to minor neurites (polarity index 36 h: 0.7 ± 0.1, 48 h: 1.7 ± 0.2; Fig 1D). While the length difference kept increasing after 36 h (48 h: 56.8 µm ± 4.6 µm; 60 h: 91.9 µm ± 4.0 µm; 72 h: 149 µm ± 7.1 µm; Fig 1D), the polarity index for acetylated tubulin (60 h: 2.2 ± 0.2; 72 h: 2.1 ± 0.2) and VAMP4 (60 h: 2.3 ± 0.3; 72 h: 1.9 ± 0.3) remained constant after 48 h (Fig 1C and D). Thus, the first indication of neuronal polarization is the elongation of a neurite relative to the others. Subsequently, the axonal marker acetylated tubulin and VAMP4-positive vesicles are polarized to the longest neurite.

**Figure 1.**
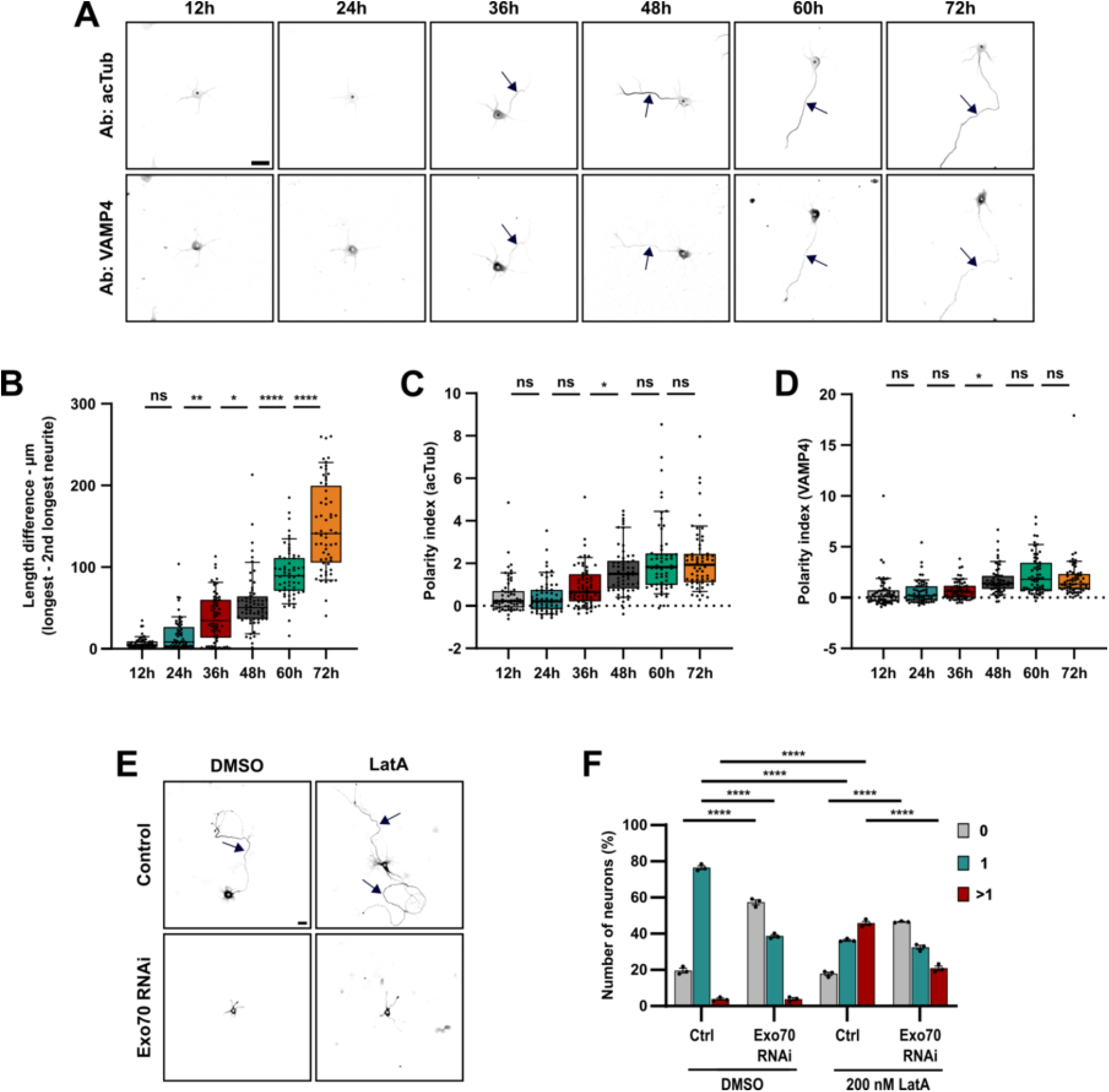
PPVs accumulate in the longest neurite. (**A**) Cortical neurons were analyzed after 12, 24, 36, 48, 60, and 72 h of culture by staining with anti-acetylated tubulin antibody (axons) and anti-VAMP4 antibodies (PPVs). (**B**) The difference in the length between the longest and the second longest neurite was quantified for (A). (**C**) The polarity index (ratio of fluorescence intensity in the longest and the second longest neurite) for acetylated tubulin (acTub) was determined for (A). (**D**) The polarity index for VAMP4 was determined for (A) (n = 3 independent experiments that are biological replicates, N = 60 neurons). (**E**) Hippocampal neurons were transfected with control (Ctrl) or knockdown vectors for Exo70 expressing EmGFP (in figure). 200 nM LatrunculinA (LatA) was added at 1 d.i.v. and axon extension analyzed at 3 d.i.v. with anti-acetylated tubulin and anti-MAP2 antibodies (minor neurites). (**F**) The percentage of neurons without an axon (0), with a single axon (1) and with multiple axons (>1) was quantified for (E) (n = 3 independent experiments that are biological replicates, N = 192 neurons). Scale bars: 20 µm. Values are means ± SEM. (**B, C, D**): one-way ANOVA, (**F**): two-way ANOVA.

The polarized distribution of PPVs is consistent with an increased demand for membrane material to allow the extension of the axon (Deng *et al*, 2014). Membrane expansion in the growing axon depends on the exocyst-dependent exocytosis of PPVs (Dupraz *et al*, 2009; Ravindran & Püschel, 2023). Axon extension in cultured neurons is limited by actin arcs in the growth cone that restrict microtubule elongation (Schelski & Bradke, 2017; Dupraz *et al*, 2019). Actin depolymerization removes this restriction and induces the formation of supernumerary axons (Bradke & Dotti, 1999; Dupraz *et al*, 2019; Lin *et al*, 2026). To investigate if this effect also depends on the exocytosis of PPVs, we tested if a knockdown of Exo70 is able to block the induction of axons after actin depolymerization by Latrunculin A (LatA). Neurons were transfected with a knockdown vector for Exo70, 200 nM LatA was added at 1 day in vitro (1 d.i.v.) and axon extension analyzed at 3 d.i.v. (Fig 1E and F, Fig S1). Depolymerization of F-actin increased the percentage of neurons with supernumerary axons from 3.8 ± 0.6% in controls to 45.8% ± 1.2%. The knockdown of Exo70 largely blocked the effect of LatA and reduced the number of neurons with multiple axons (control + LatA: 45.8 ± 1.2% neurons with multiple axons; Exo70 RNAi + LatA: 20.9 ± 1.3%; control + LatA: 17.9 ± 0.9% neurons without an axon; Exo70 RNAi + LatA: 46.6 ± 0.2%; Exo70 RNAi: 57.4 ± 1.4%). These results suggest that the Exo70-dependent exocytosis of PPVs is required for axon extension even when the restrictions imposed by the cytoskeleton are removed, highlighting the importance of membrane expansion.

### PPV distribution depends on neurite length

Previous work has shown that a neurite is selected as the axon when it has grown sufficiently to establish a critical difference in length relative to the other neurites (Goslin & Banker, 1989; Yamamoto *et al*, 2012). Our results confirmed that the first indication for the establishment of an asymmetry is a persistent length difference. To determine if this difference is also required for the polarization of PPVs to the nascent axon, we generated microstructures that either restrict neurite extension to 30 µm (R structure) or allow the extension of one neurite to 120 µm (UnR structure). In these microstructures, a rhomboid center that allows the attachment of the soma is connected to two perpendicular arms (Fig 2A). The short length of these arms in R structures is expected to prevent the formation of an axon by restricting neurite extension while the single long arm allows the extension of an axon in UnR structures (Yamamoto *et al*, 2012). To determine how a restriction of neurite extension affects the distribution of PPVs, we quantified the polarity index for VAMP4 in unpolarized neurons (R) and neurons with a single axon (UnR) at 48 h (Fig 2A). The majority of neurons (88.2 ± 5.9%; Fig 2B) did not extend an axon in R structures (polarity index for acetylated tubulin in R: 0.1 ± 0.1, UnR: 2.3 ± 0.5; Fig 2D) while most neurons extended a single axon in UnR structures (85.2 ± 8.3%), consistent with previous results (Yamamoto *et al*, 2012). VAMP4 becomes preferentially localized to a single neurite only when one neurite can elongate while restricting extension prevents it (polarity index UnR: 1.3 ± 0.4, R: 0.3 ± 0.2; UnR: 66.4 ± 5.7 µm length difference, R: 9.6 ± 1.7 µm; Fig 2C - 2E). These results suggest that the establishment of a difference in neurite length is an important step in neuronal polarization that is followed by an accumulation of PPVs in a single neurite and axon extension.

**Figure 2.**
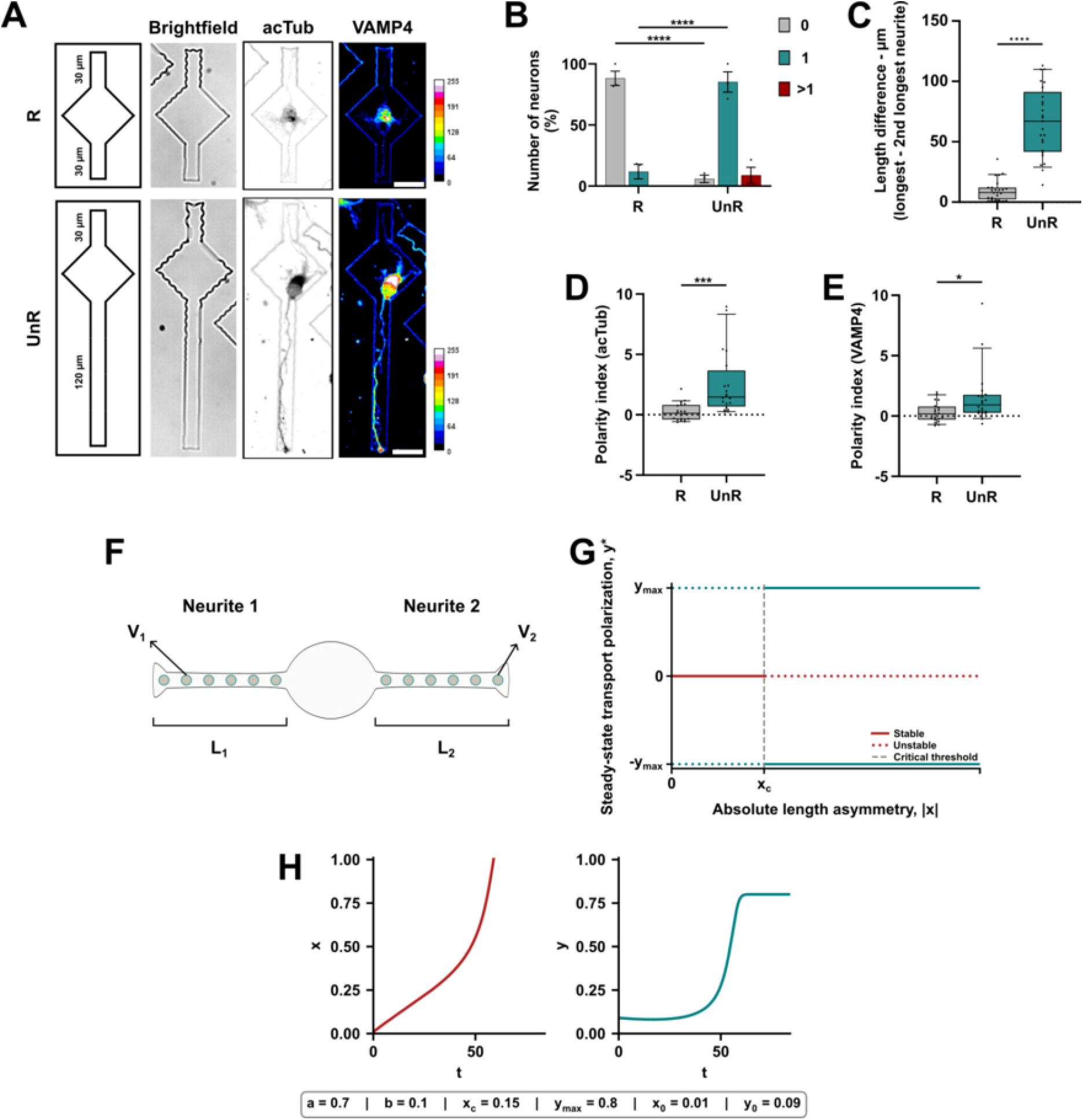
PPV distribution depends on neurite length. **(A)** Schematic representations and bright field images of the microstructures used for culturing neurons are shown. A rhomboid center for the soma is connected to two perpendicular arms of 30 µm (restricted, R) or two arms of 30 and 120 µm (unrestricted, UnR). Cortical neurons were cultured in dishes with R or UnR microstructures and analyzed at 2 d.i.v. (48 h) by staining with anti-acetylated tubulin antibody (axons) and anti-VAMP4 antibodies (PPVs). (**B**) The percentage of neurons without an axon (0), with a single axon (1) and with multiple axons (>1) was quantified for (A). (**C**) The difference in the length between the longest and the second longest neurite was quantified for (A). (**D**) The polarity index for acetylated tubulin was determined for (A). (**E**) The polarity index for VAMP4 was determined for (A) (n = 3 independent experiments that are biological replicates, N = 25 neurons). (**F**) Schematic illustration of the model for a length-dependent polarization of vesicle transport. The model assumes two competing neurites (neurite 1 and 2) with the length *L*_1_(*t*) and *L*_2_(*t*), respectively. A common pool supplies vesicles that are transported into both neurites at rates *V*_1_(*t*) and *V*_2_(*t*). **(G**) The diagram shows the steady-state transport polarization (*y*^∗^) as a function of the absolute normalized neurite length asymmetry (| *x* |). Below the critical threshold *x*_c_, the unpolarized state (*y*^∗^ = 0) is stable, whereas the polarized states (*y*^∗^ = ±*y*_max_) are unstable. Above *x*_c_, the unpolarized state becomes unstable, while the two symmetry-related polarized states become stable. Solid lines indicate stable steady states, and dotted lines indicate unstable steady states. The vertical dashed line marks the critical threshold *x*_c_, at which the system becomes degenerate and all polarization values within the allowed range are steady states. The diagram illustrates the threshold-dependent stability exchange predicted by the model. (**H**) Solutions *x*(*t*) and *y*(*t*) for the numerical integration of the coupled dynamical system, 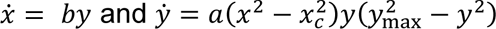, obtained for parameters as indicated in the graph. The length asymmetry diverges whereas transport polarization reaches saturation after a certain time. Arrows mark axons. Scale bars: 20 µm. Values are means ± SEM. (**B**): two-way ANOVA, (**C, D, E)**: unpaired t test.

To investigate conditions that allow an initiation of a polarized vesicle transport by the emergence of a length asymmetry that is suggested by the biased VAMP4 localization, we developed a mathematical framework that links neurite length and vesicle transport (Supplementary Note 1, Fixed-length transport model). In this framework, the linear stability of transport allocation depends on the normalized length difference between neurites. We modelled two initially equivalent neurites (Fig 2F) using their normalized length asymmetry (*x*) and a variable describing the polarization of PPV transport (*y*) between them. The resulting bifurcation diagram depicts the linear stability of the steady states of the dynamical system (Fig 2G). It predicts a critical length asymmetry below which small transport biases decay and above which the unpolarized symmetric transport state becomes unstable and polarized states stable (Strogatz, 2018). This strongly simplified framework gives a potential explanation for why polarized VAMP4 localization may emerge under growth-permissive conditions but would remain absent when neurite extension is restricted.

The fixed-length transport model does not account for the evolution of differences in neurite length over time. We extended this model to a coupled dynamical system to better delineate the connection between morphological asymmetry and the polarization of vesicle transport (Supplementary Note 1). Given that a preferential vesicle delivery may promote neurite elongation through membrane insertion, we introduced a minimal feedback that links vesicle transport and neurite growth (Supplementary Note 1, Minimal coupled dynamical system). Numerical simulations were used to investigate how neurite length asymmetry *x*(*t*) and polarization of vesicle transport *y*(*t*) develop over time (Fig S2).

Variation of different parameters reveals that the polarization of transport emerges from a competition between the decay of a small transport bias below the critical threshold and its amplification once the threshold is crossed. Within a specific parameter range, the change of length asymmetry and transport polarization over time are comparable to those observed in cultured neurons, as length asymmetry diverges and transport polarization saturates (Fig 2H). In particular, these simulations indicate that a strong coupling of neurite length with polarized transport and membrane insertion can mediate an amplification of an initial, small morphological asymmetry and facilitate the entry into the polarized regime.

These theoretical considerations propose a length-dependent polarization mechanism that depends on vesicle transport and exocytosis, and suggest experimental tests to establish a connection between these processes.

### Increasing the supply of PPVs promotes axon formation

The accumulation of PPVs in the longest neurite during neuronal polarization raises the question if an increased transport of PPVs is sufficient to promote axon formation. VAMP4-positive vesicles are anterogradely transported into the axon by Kinesin-1 (Deng *et al*, 2014). To determine whether an increase in vesicle supply is sufficient to promote axon formation, we implemented a heterodimerizer system to recruit constitutively active forms of kinesins that mediate the transport of various axonal cargos (Kif1A_ ca-DmrC or Kif5C_ca-DmrC) to PPVs using the PPV marker Rab10 (DmrA-Rab10; Fig 3A) (Jacobson *et al*, 2006; Wang *et al*, 2011; Jenkins *et al*, 2012; Deng *et al*, 2014; van Bergeijk *et al*, 2015). Addition of the heterodimerizer to neurons transfected with these constructs leads to an increase in vesicle supply by their anterograde transport (Kapitein *et al*, 2010; Jenkins *et al*, 2012; van Bergeijk *et al*, 2015). Stimulating the transport of PPVs by linking them to either active Kif1A or Kif5C by addition of dimerizer at 22 h induced the extension of supernumerary axons (Fig 3B and C, Fig S3A). The percentage of neurons with multiple axons increased from 15.3 ± 1.2% to 38.4 ± 0.7% (Kif1A) and from 13.2 ± 3.5% to 31.1 ± 1.4% (Kif5C), respectively. The percentage of neurons with multiple axons did not change significantly in controls (DmrC + solvent: 15.1 ± 0.3%, DmrC + dimerizer: 12.6 ± 1.2%).

**Figure 3.**
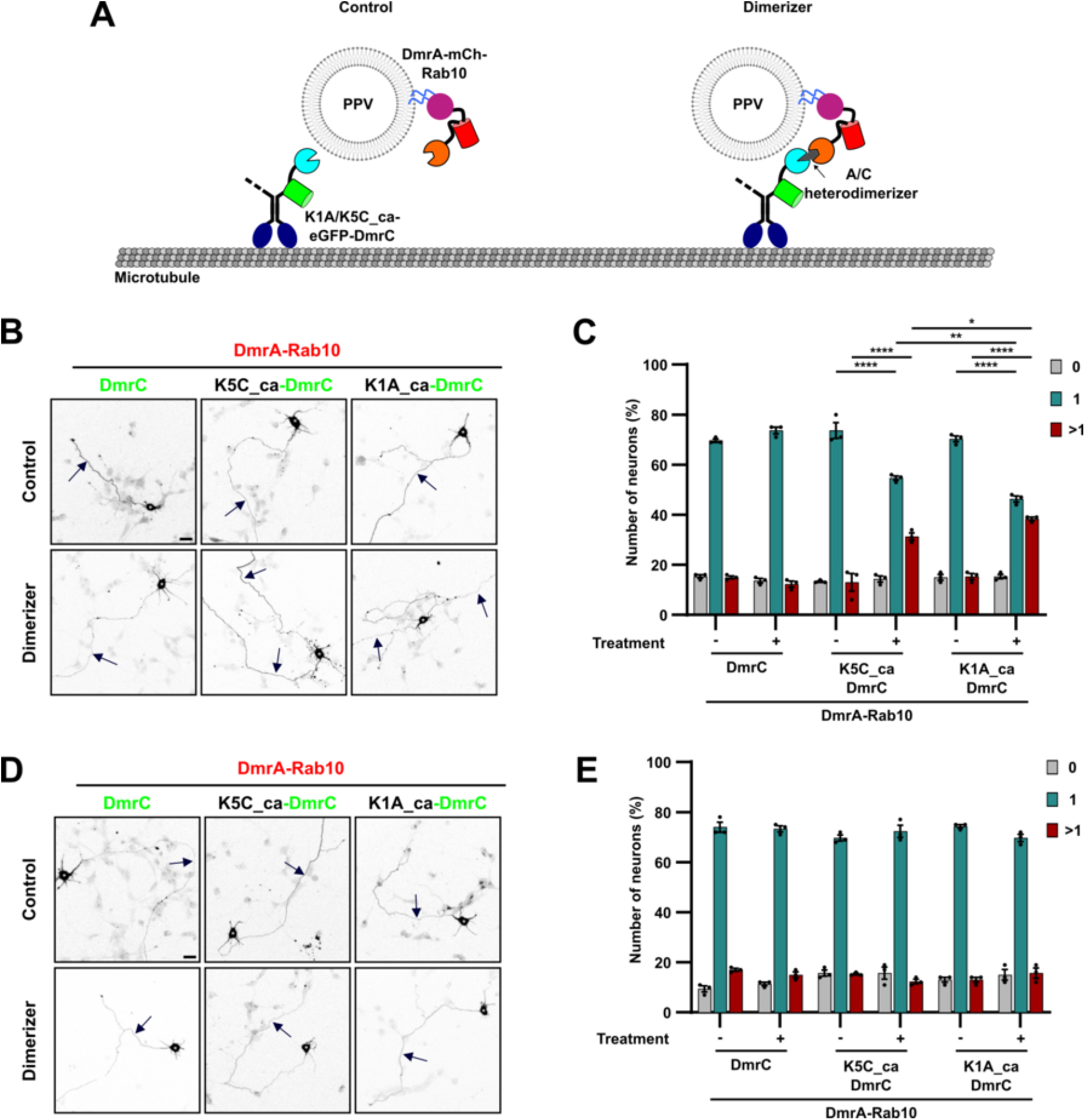
Increased PPV transport induces the extension of supernumerary axons. (**A**) Schematic representation of the inducible heterodimerization system used to increase the transport of PPVs into neurites. Fusion proteins of constitutively active (ca) Kif5C or Kif1A with DmrC and eGFP (K1A/K5C_ca-eGFP-DmrC), and DmrA with mCherry and Rab10 (DmrA-mCh-Rab10) were expressed in neurons. EGFP-DmrC without a motor domain served as negative control. To recruit active Kinesins to PPVs, the A/C heterodimerizer was added. (**B**, **D**) Cortical neurons were co-transfected with vectors for DmrC (control), K1A_ca-DmrC or K5C_ca-DmrC, DmrA-mCh-Rab10 and sfGFP to label transfected neurons (shown in B and D). Solvent (DMSO, control) or heterodimerizer (0.25 nM) was added at 22 or 34 h of culture for 30 minutes. Axon extension was analyzed at 3 d.i.v. with anti-acetylated tubulin and anti-MAP2 antibodies. (**C**, **E**) The percentage of neurons without an axon (0), with a single axon (1) and with multiple axons (>1) was quantified for (B) and (D) (n = 3 independent experiments that are biological replicates, N > 300 neurons). Arrows mark axons. Scale bars: 20 µm. Values are means ± SEM. **(C, E):** two-way ANOVA.

When dimerizer was added at 34 h, after most neurons already extended an axon and the transport of axonal cargo has become selective, no effect was observed (Kif5C-DmrC/control: 15.1 ± 0.5%, Kif5C-DmrC /dimerizer: 12.0 ± 0.9%; Kif1A-DmrC /control: 12.7 ± 1.1%, Kif1A-DmrC/dimerizer: 15.7 ± 2.1%; DmrC/control: 16.8 ± 0.7% to 15.1 ± 1.2% DmrC/dimerizer; Fig 3D and E, Fig S3B). These results indicate that increasing the transport of PPVs to neurites promotes axon formation when vesicles are supplied before polarization is established.

### Rab10 activation on PPVs promotes axon formation

Since an increased PPV transport promoted axon extension, we next analyzed the role of Rab10 that is involved in both, the transport and exocytosis of PPVs (Wang *et al*, 2011; Deng *et al*, 2014). We tested whether Rab10 activation on PPVs or at the plasma membrane was sufficient to promote axon extension. Previous studies showed that recruiting the catalytic DENN domain of the Rab10 GEF DENND2B activates Rab10 on the outer mitochondrial membrane (Kumar *et al*, 2022). We adapted this technique to recruit the DENN domain (eGFP-DENND(2B)-DmrC) to PPVs by heterodimerization with DmrA-mCh-Rab10 or DmrA-mCh-VAMP4, or to the plasma membrane using a myristoylation sequence (myr-mCh-DmrA) (Fig 4A). Solvent (control) or dimerizer were added to transfected neurons at 22 h of culture before neuronal polarization, and axon extension was analyzed at 3 d.i.v. (Fig 4B and 4C, Fig S4). Recruitment of DENN(2B)-DmrC to PPVs by either DmrA-Rab10 (control: 16.6 ± 0.4%, dimerizer: 38.2 ± 1.7%) or DmrA-VAMP4 (control: 17.9 ± 1.4%, dimerizer: 37.6 ± 1.5%) induced the extension of supernumerary axons (Fig 4B, C). When DENND(2B)-DmrC was directed to the plasma membrane, the percentage of neurons with multiple axons did not differ significantly from control (control: 18.6 ± 3.1%, dimerizer: 17.8 ± 0.5%). These results indicate that the activation of Rab10 on PPVs is sufficient to promote axon extension. In contrast, activation at the plasma membrane does not have an effect.

**Figure 4.**
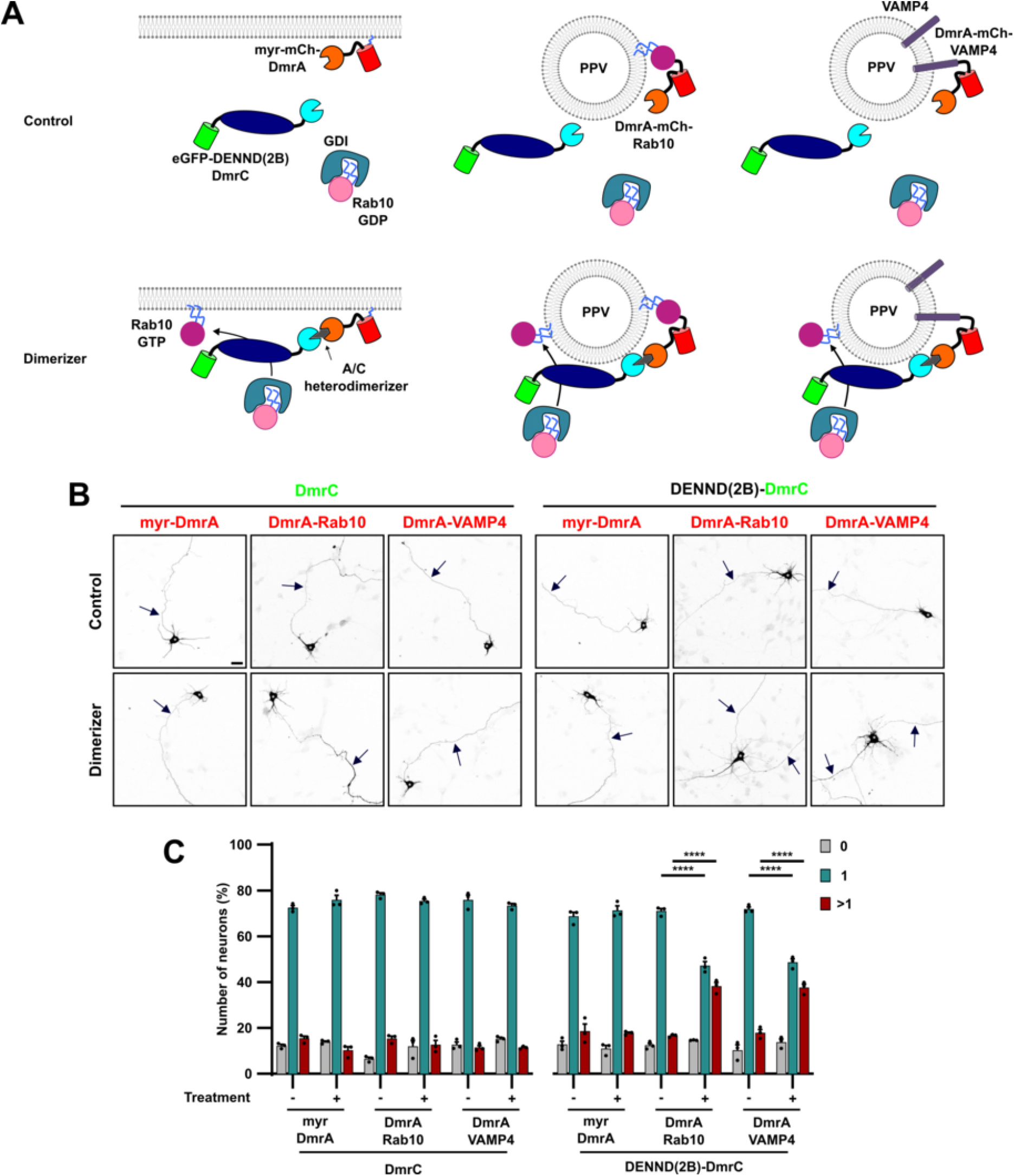
Rab10 activation on PPVs promotes axon formation. (A) Schematic representation of the inducible heterodimerization system used to activate Rab10. The catalytic DENN domain of DENND2B was fused to DmrC and eGFP (eGFP-DENND(2B)-DmrC) to activate Rab10 on membranes. To recruit it to PPVs, fusion proteins of DmrA, mCherry, and Rab10 (DmrA-mCh-Rab10) or VAMP4 were used (DmrA-mCherry-VAMP4). To target Rab10 activation to the plasma membrane, DmrA was linked to a myristoylation sequence (myr-mCh-DmrA). EGFP-DmrC without a DENN domain served as negative control. DmrC-fusion proteins are recruited by adding the A/C heterodimerizer. (B) Cortical neurons were co-transfected with vectors for DmrC (control) or DENND(2B)-DmrC, DmrA-Rab10, DmrA-VAMP4 or myr-DmrA, and sfGFP to label transfected neurons (shown in B and D). Solvent (DMSO, control) or heterodimerizer was added for 30 minutes at 22 h of culture. Axon extension was analyzed at 3 d.i.v. with anti-acetylated tubulin and anti-MAP2 antibodies. (**C**) The percentage of neurons without an axon (0), with a single axon (1) and with multiple axons (>1) was quantified for (B) (n = 3 independent experiments that are biological replicates, N > 300 neurons). Values are means ± SEM. Arrows mark axons. Scale bars: 20 µm. **(C)**: two-way ANOVA.

### Active Arhgef7 promotes axon extension through Cdc42b

Previous studies showed that Arhgef7 promotes the exocytosis of PPVs and extension of axons by activating the GTPases Cdc42b and TC10 upstream of the exocyst subunit Exo70 (Dupraz *et al*, 2009; López Tobón *et al*, 2018; Ravindran & Püschel, 2023). We generated constitutively active forms of Arhgef7 by introducing two mutations (W200K and Y599E; Fig 5A) (Marston *et al*, 2020). The constitutive activity of these Arhgef7 mutants was confirmed by their ability to induce supernumerary axons (Fig S5A and B). To show that the constitutively active Arhgef7 W200K,Y599E mutant (Arhgef7-ca) is able to activate the brain-specific isoform of Cdc42 (Cdc42b), we expressed wild type Arhgef7 or active Arhgef7-ca together with Cdc42b in HEK 293T cells and quantified the amount of GTP-bound Cdc42b using a GST-pulldown assay with GST-CRIB (Bagrodia *et al*, 1998; Neudauer *et al*, 1998). While wild type Arhgef7 increased the amount of active Cdc42b slightly (relative activity in control: 1; Arhgef7: 4.8 ± 1.8), Arhgef7-ca showed a significantly increased activity (30.2 ± 14.7; Fig 5B and C).

**Figure 5.**
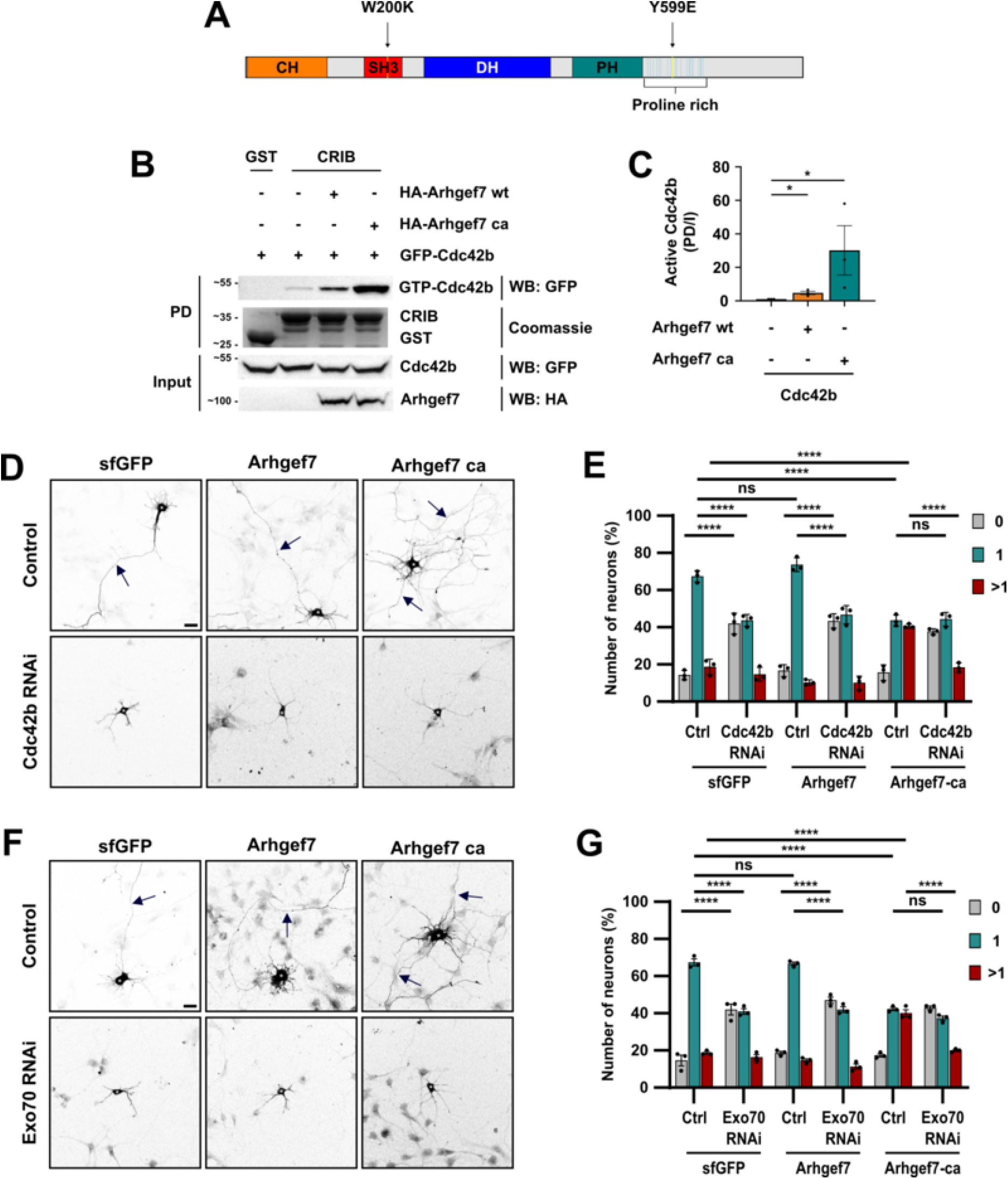
Arhgef7 acts upstream of Cdc42b and Exo70 to promote axon extension. (**A**) The mutations introduced into the SH3 domain and the proline-rich region of Arhgef7 to create constitutively active mutants (ca: W200K,Y599E) are indicated by arrows. (**B**) HA-Arhgef7 or HA-Arhgef7-ca and eGFP-Cdc42b were expressed in HEK 293T cells and the amount of active GTP-Cdc42 determined using a pulldown assay (PD) with GST-CRIB, and analyzed by Western blot (WB) or Coomassie Blue staining. The molecular weight is indicated in kDa. (**C**) The amount of active Cdc42b was quantified for (B). The relative intensity of the control experiment was defined as 1 (n=3 independent experiments that are biological replicates). (**D**) Hippocampal neurons were co-transfected with control (Ctrl) or RNAi vectors for Cdc42b expressing H2B-RFP and vectors for sfGFP, sfGFP-Arhgef7 or sfGFP-Arhgef7-ca (sfGFP shown in D). Axon extension was analyzed at 3 d.i.v. with anti-acetylated tubulin and anti-MAP2 antibodies. (**E**) The percentage of neurons without an axon (0), with a single axon (1) and with multiple axons (>1) was quantified for (D) (n = 3 independent experiments that are biological replicates, N > 300 neurons). (**F**) Hippocampal neurons were co-transfected with control (Ctrl) or RNAi vectors for Exo70 expressing H2B-RFP and vectors for sfGFP, sfGFP-Arhgef7 or sfGFP-Arhgef7-ca (sfGFP shown in F). Axon extension was analyzed at 3 d.i.v. with anti-acetylated tubulin and anti-MAP2 antibodies. (**G**) The percentage of neurons without an axon (0), with a single axon (1) and with multiple axons (>1) was quantified for (F) (n = 3 independent experiments that are biological replicates, N > 300 neurons). Arrows mark axons. Scale bars: 20 µm. Values are means ± SEM. **(C)**: ratio paired Student’s t-test, **(E, G):** two-way ANOVA. Source data are available for this figure.

To confirm that it can activate Cdc42b also in neurons, we analyzed axon extension at 3 d.i.v. after expression of Arhgef7-ca together with a Cdc42b knockdown construct (Fig 5D and E, Fig S5C) (Ravindran & Püschel, 2023). While wild type Arhgef7 did not have an effect, Arhgef7-ca induced the formation of supernumerary axons (wt: 10.1 ± 0.9%, ca: 40.6 ± 0.8%), which was blocked by the knockdown of Cdc42b (control: 14.2 ± 1.4% neurons without axons; Cdc42b RNAi: 42.0 ± 3.3%; Arhgef7-ca + Cdc42b RNAi: 37.6 ± 0.7%; Arhgef7-ca + Cdc42b RNAi: 18.2 ± 1.7% neurons with multiple axons; Fig 5E). To test if Arhgef7-ca also acts upstream of Exo70, we expressed Arhgef7-ca together with a Exo70 knockdown construct (Ravindran & Püschel, 2023). Similar to the effect of the Cdc42b RNAi construct, the Exo70 knockdown blocked axon extension in control neurons and in neurons expressing Arhgef7-ca (control: 14.4 ± 2.8% neurons without an axon, Exo70 RNAi: 42.1 ± 3.0%; Arhgef7-ca + Exo70 RNAi: 42.8 ± 0.8%; Arhgef7-ca: 40.2 ± 1.7% neurons with multiple axons; Arhgef7-ca + Exo70 RNAi 19.9 ± 0.7%; Fig 5F and G, Fig S5D). These results confirm that Arhgef7-ca promotes axon extension upstream of Cdc42b and Exo70 (Ravindran & Püschel, 2023).

### Rab10 interacts with and activates Arhgef7

Since both, Rab10 and Arhgef7, promote exocytosis and axon extension (Wang *et al*, 2011; Deng *et al*, 2014; López Tobón *et al*, 2018; Ravindran & Püschel, 2023), we tested whether they interact. A pulldown experiment showed a clear interaction of active GST-Rab10-S18V and Arhgef7 after expression in HEK-293T cells (Fig 6A). Interestingly, no interaction was detectable with the Arhgef7-ca mutant, suggesting that Rab10 binds only to the inactive form of Arhgef7. To test whether Rab10 stimulates Arhgef7 activity, constitutively active Rab10 S18V (Rab10-ca) was expressed together with wild type Arhgef7 and Cdc42b in HEK 293T cells and the amount of active Cdc42b-GTP quantified by a pull-down with GST-CRIB (Fig 6B and C). The co-expression of Rab10-ca significantly stimulated the activity of Arhgef7 and increased the amount of active Cdc42 (relative activity in control: 1; Arhgef7: 1.9 ± 0.6; Arhgef7 + Rab10-ca: 70.3 ± 20.3). These results show that Rab10 not only interacts with Arhgef7 but also stimulates its activity as a GEF for Cdc42b.

**Figure 6.**
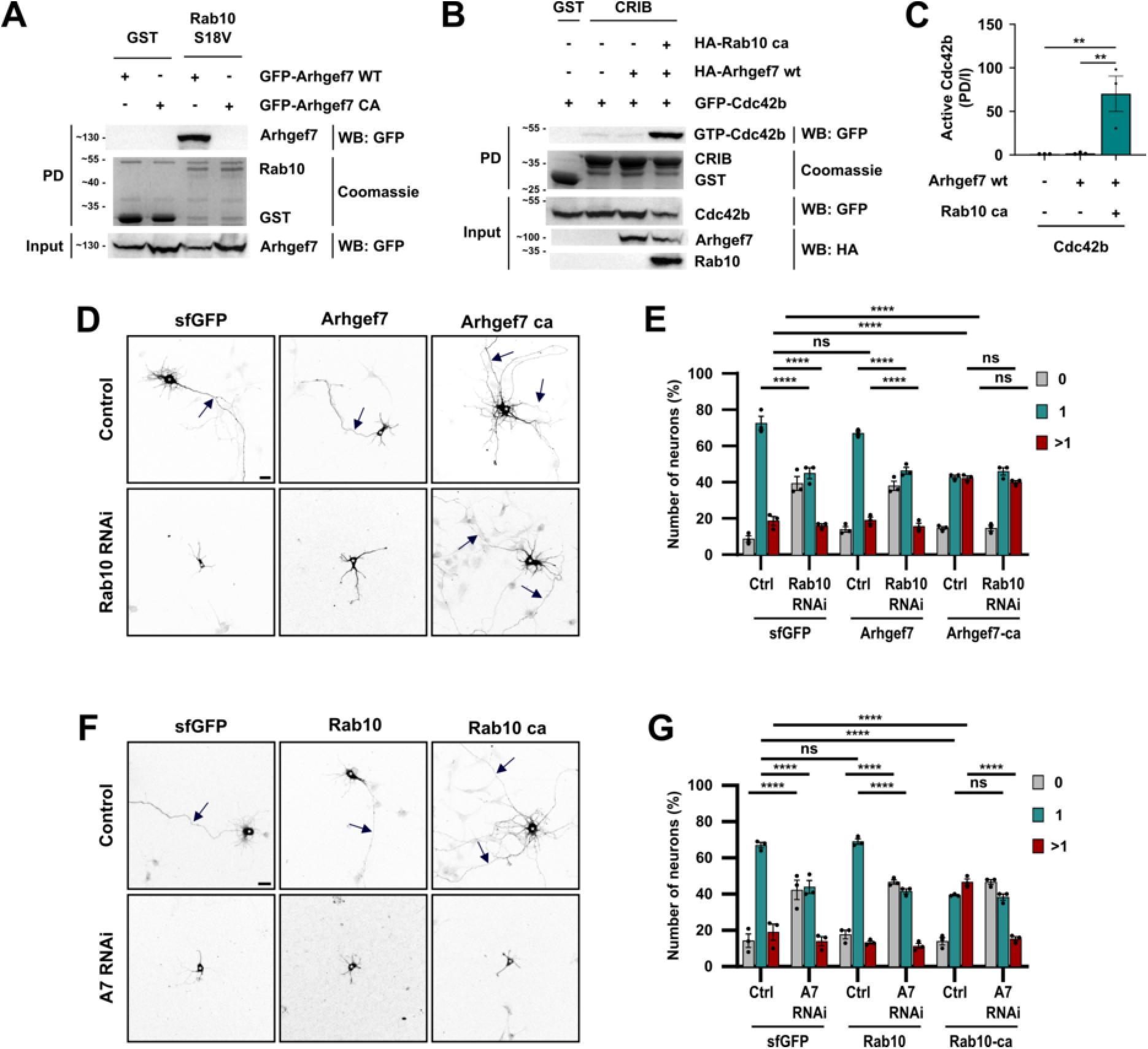
Rab10 interacts with and activates Arhgef7. (**A**) eGFP-Arhgef7 or eGFP-Arhgef7-ca were expressed in HEK 293T cells and the interaction with Rab10 tested using a pulldown assay with GST (control) or GST-Rab10-S18V (ca) (n=3 independent experiments that are biological replicates). (**B**) HA-Rab10-S18V (ca) was co-expressed with HA-Arhgef7 or HA-Arhgef7 and eGFP-Cdc42b in HEK293T cells as indicated. The amount of active GTP-Cdc42 was determined by a pulldown assay (PD) with GST-CRIB and Coomassie Blue staining, and analyzed by Western blot (WB). The molecular weight is indicated in kDa. (**C**) The amount of active Cdc42b was quantified for (B) (n=3 independent experiments that are biological replicates). (**D**) Hippocampal neurons were co-transfected with control (Ctrl) or RNAi vectors for Rab10 expressing emGFP and vectors for sfGFP, sfGFP-Arhgef7 or sfGFP-Arhgef7-ca (shown in D). Axon extension was analyzed at 3 d.i.v. with anti-acetylated tubulin and anti-MAP2 antibodies. (**E**) The percentage of neurons without an axon (0), with a single axon (1), and with multiple axons (>1) was quantified for (D) (n = 3 independent experiments that are biological replicates, N > 300 neurons). (**F**) Hippocampal neurons were co-transfected with control (Ctrl) or RNAi vectors for Arhgef7 expressing H2B-RFP and vectors for sfGFP, sfGFP-Rab10 or sfGFP-Rab10-ca (shown in G). Axon extension was analyzed at 3 d.i.v. with anti-acetylated tubulin and anti-MAP2 antibodies. (**G**) The percentage of neurons without an axon (0), with a single axon (1) and with multiple axons (>1) was quantified for (F) (n = 3 independent experiments that are biological replicates, N > 300 neurons). Arrows mark axons. Scale bars: 20 µm. Values are means ± SEM. **(C):** ratio paired Student’s t-test, **(E, G)**: two-way ANOVA. Source data are available for this figure.

To address the question if Rab10 acts upstream of Arhgef7 in neurons, we first tested if the expression of Arhgef7 or Arhgef7-ca can rescue the knockdown Rab10 (Fig 6D, E). Consistent with previous results (Wang *et al*, 2011), the knockdown of Rab10 impairs axon extension, which can be rescued by an RNAi-resistant Rab10 construct, confirming the specificity of the knockdown (Fig S6A-D). The percentage of unpolarized neurons without an axon increased from 8.7 ± 1.8% in controls to 38.0% ± 2.5% after knockdown of Rab10 (Fig 6D and E). The expression of wild type Arhgef7 could not rescue the suppression of Rab10 while Arhgef7-ca was able to counteract the loss of Rab10. Arhgef7-ca decreased the number of unpolarized neurons from 39.0 ± 3.8% to 14.3 ± 1.5% after Rab10 knockdown and induced the formation of supernumerary axons (control + sfGFP: 18.8 ± 2.3% neurons with multiple axons; control + Arhgef7-ca: 42.4 ± 1.1%; Rab10 RNAi + Arhgef7-ca: 39.9 ± 1.0%; Fig 6D and E, Fig S6E). These results indicate that Arhgef7 acts downstream of Rab10. To investigate if Rab10 also requires Arhgef7 to promote axon extension, we tested if the knockdown of Arhgef7 blocks the induction of supernumerary axons by active Rab10 Q68L (Rab10-ca) (Wang *et al*, 2011; Ravindran & Püschel, 2023). The expression of Rab10-ca but not wild type Rab10 increased the number of neurons with multiple axons (control (sfGFP): 19.1% ± 4.3%, Rab10-ca: 47.0 ± 1.9, Rab10: 13.2 ± 1.0%; Fig 6F and G, Fig S6F), which was blocked by the knockdown of Arhgef7 (Rab10 + Arhgef7 RNAi: 46.8 ± 1.0% neurons without an axon; Rab10-ca + Arhgef7 RNAi: 46.4 ± 1.2%; sfGFP + Arhgef7 RNAi: 41.9 ± 5.3%). These results show that the interaction with Rab10 activates Arhgef7 and promotes axon extension.

### Recruitment of Arhgef7 to PPVs promotes axon extension

Since Arhgef7 plays a critical role in PPV exocytosis (Ravindran & Püschel, 2023) and its activity is regulated by Rab10, it may link the transport of PPVs and their subsequent exocytosis. In this case, an activation of Arhgef7 on PPVs should be sufficient to promote axon extension. To test if Arhgef7 activation is required on PPVs or at the plasma membrane, we recruited the catalytic DHPH domain of Arhgef7 (Arhgef7_DHPH-DmrC) to PPVs (DmrA-Rab10, DmrA-VAMP4) or the plasma membrane (myr-DmrA). EGFP-DmrC without the catalytic domain served as negative control. The catalytic DHPH1 domain of the Cdc42 GEF Fgd1 was used as a positive control for Cdc42 activation at the plasma membrane (Woo *et al*, 2019). Dimerizer was added to transfected neurons at 22 h of culture, before axons are formed, and axon extension analyzed at 3 d.i.v. (Fig 7, Fig S7). The Arhgef7 DHPH domain did not have a significant effect on axon formation when recruited to the plasma membrane (control: 14.3% ± 0.2% neurons with multiple axons; dimerizer: 11.0 ± 1.5%). By contrast, Fgd1 DHPH1 localization to the plasma membrane promoted the extension of supernumerary axons (myr-DmrA control: 14.4 ± 1.1% neurons with multiple axons, dimerizer: 36.4 ± 0.5%) as reported before (Woo *et al*, 2019).

**Figure 7.**
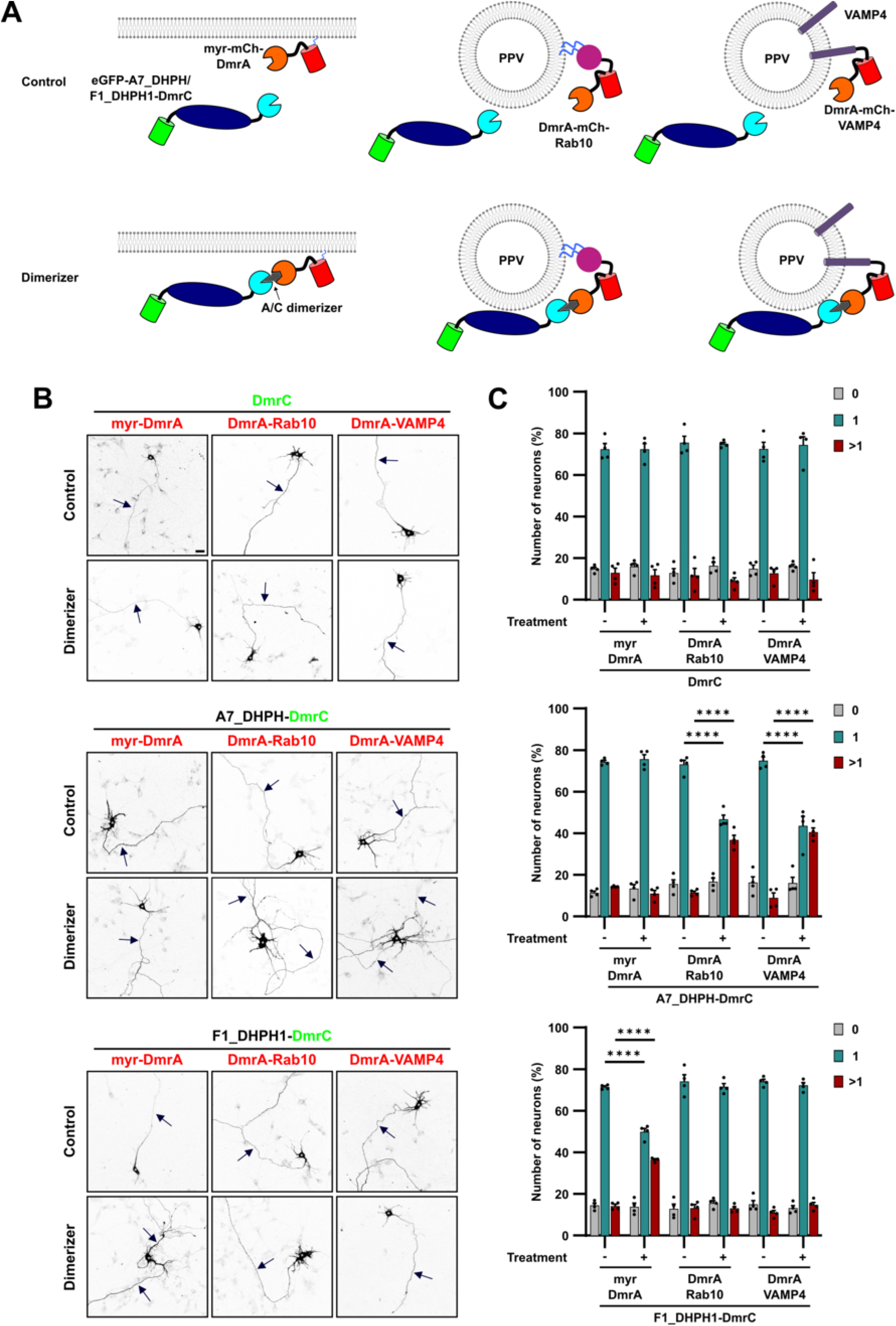
Recruitment of Arhgef7 to PPVs promotes axon extension. (**A**) The catalytic DHPH domains of Arhgef7 or Fgd1 were fused to DmrC and eGFP (eGFP-A7_DHPH-DmrC, eGFP-F1_DHPH1-DmrC). DmrA with a myristoylation sequence (myr-mCh-DmrA) was used to target DHPH domains to the plasma membrane. Fusion proteins of DmrA with mCherry and Rab10 or VAMP4 (DmrA-mCh-Rab10, DmrA-mCherry-VAMP4) were used for recruitment to PPVs. EGFP-DmrC without a catalytic domain served as negative control. (**B**) Cortical neurons were co-transfected with vectors for eGFP-A7_DHPH-DmrC, eGFP-F1_DHPH1-DmrC or DmrC, myr-DmrA, DmrA-Rab10 or DmrA-VAMP4 and sfGFP to label transfected cells (sfGFP shown in B). Vehicle (DMSO, control) or dimerizer was added at 22 h of culture for 30 min. Axon extension was analyzed at 3 d.i.v. with anti-acetylated tubulin and anti-MAP2 antibodies. (**C**) The percentage of neurons without an axon (0), with a single axon (1) and with multiple axons (>1) was quantified for (B) (n = 4 independent experiments that are biological replicates, N > 400 neurons). Arrows mark axons. Scale bars: 20 µm. Values are means ± SEM. **(C)**: two-way ANOVA.

However, recruitment of Fgd1 DHPH1 to PPVs did not have an effect (DmrA-Rab10 control: 13.1% ± 1.8% neurons with multiple axons, DmrA-Rab10 + dimerizer: 13.0% ± 1.0%; DmrA-VAMP4 control: 12.4 ± 1.6%, DmrA-VAMP4 + dimerizer: 14.7 ± 1.3%). By contrast, recruitment of Arhgef7 DHPH to PPVs significantly increased the number of neurons with multiple axons (DmrA-Rab10 control: 11.5 ± 0.8%, DmrA-Rab10 + dimerizer: 36.7 ± 2.3%; DmrA-VAMP4 control: 8.9 ± 2.4%, DmrA-VAMP4 + dimerizer: 40.5 ± 2.0%). These results show that Arhgef7 activity is required specifically on PPVs and not at the plasma membrane, suggesting that Rab10 interacts with and activates Arhgef7 on PPVs, leading to axon extension.

## Discussion

Neurons become polarized when one of their neurites begins to elongate continuously as the axon (Dotti *et al*, 1988). Axon extension requires an expansion of the plasma membrane, which depends on the polarization of transport and exocytosis of PPVs to the growing neurite (Craig *et al*, 1995; Futerman & Banker, 1996; Bradke & Dotti, 1997; Pfenninger, 2009). Here we show that increasing the transport of PPVs before neurons are polarized is sufficient to induce the extension of supernumerary axons. This effect indicates that an increased supply of vesicles is connected to their subsequent insertion into the membrane to allow neurite growth. The link between vesicle transport and exocytosis is mediated by the interaction of Rab10 with Arhgef7. Rab10 stimulates Arhgef7 activity on PPVs as a GEF for Cdc42b to promote axon extension through exocyst-dependent exocytosis (Fig 8).

**Figure 8.**
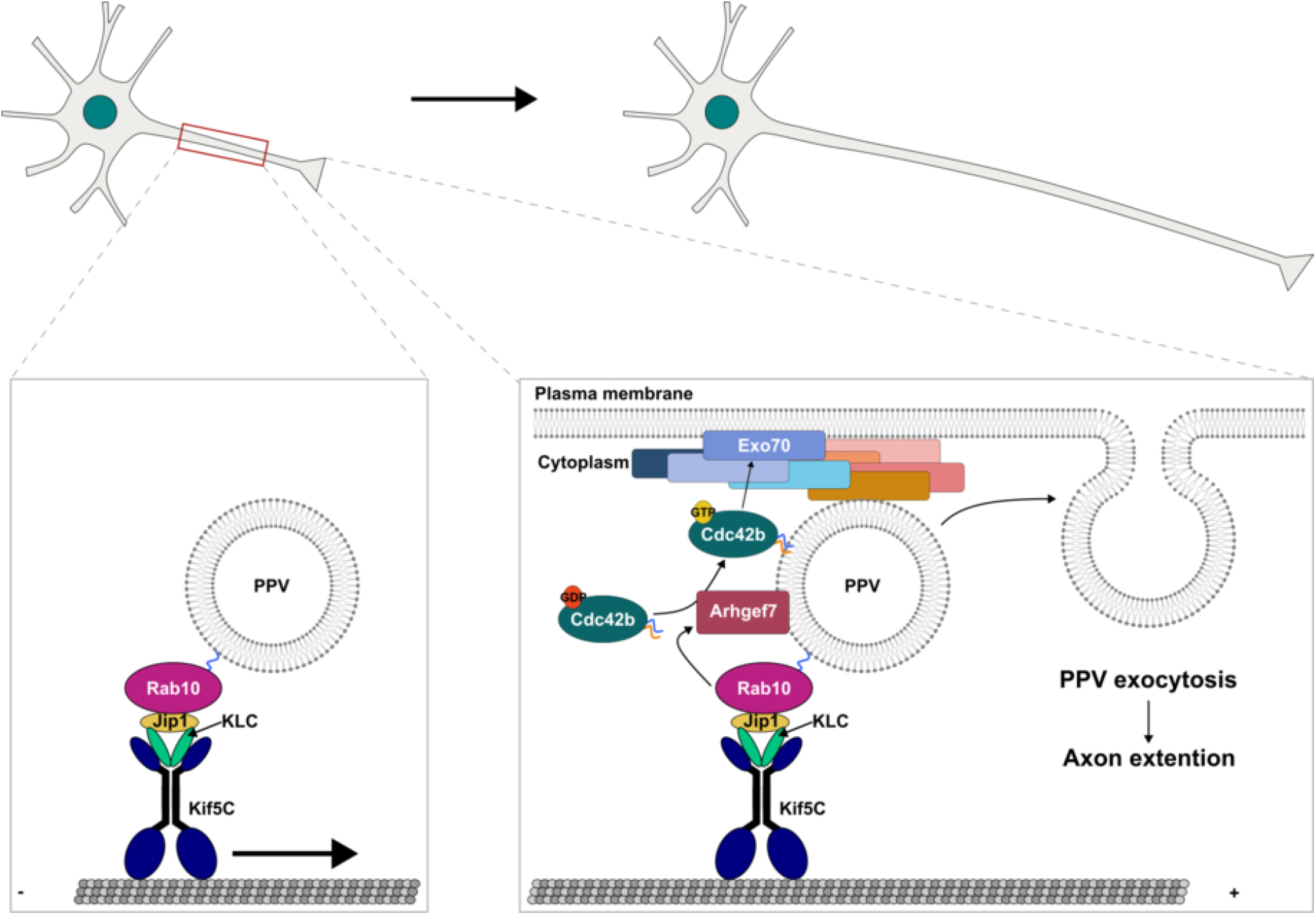
Rab10 and Arhgef7 promote axon extension by linking PPV supply and exocytosis. An asymmetry in neurite length precedes axon specification, which coincides with an accumulation of PPVs in the nascent axon. The anterograde transport of PPVs by Kinesin-1 supplies PPVs to the growth cone of extending neurites. The exocyst tethers PPVs at the plasma membrane prior to their exocytosis. In the growth cone, PPVs stimulate their insertion into the membrane to promote neurite growth. The link between vesicle transport and exocytosis is mediated by the interaction of Rab 10 and Arhgef7, which activates Cdc42b. The interaction of active Cdc42b with the exocyst subunit Exo70 promotes axon extension by increasing exocyst-dependent exocytosis. This signaling cascade links PPV transport in the growing axon to the subsequent insertion into the plasma membrane.

An analysis of immature neurons indicates that the distribution of PPVs is biased towards the longest neurite after neurons show a persistent difference in neurite length, which was the first detectable asymmetry between the neurites. The axonal marker acetylated tubulin and the PPV marker VAMP4 became polarized to the longest neurite only after a significant difference in length was detectable. To further explore the possibility that an asymmetry of neurite length and the polarization of vesicle transport interact during neuronal polarization, we formulated a mathematical framework of these processes. This phenomenological, nonlinear dynamical system describes how an existing transport bias may be amplified and stable polarized states arise under appropriate conditions. It shows that the morphologically symmetric state remains locally unstable, indicating that a critical length asymmetry threshold can trigger vesicle transport polarization from small perturbations. These perturbations could result from stochastic fluctuations in transport, the arrival of actin waves or external cues (Namba *et al*, 2015; Winans *et al*, 2016; Schelski & Bradke, 2017; Takano *et al*, 2019; Lin *et al*, 2026). The effect of an experimental restriction of neurite extension with patterned surfaces on PPV distribution supports an association between neurite length and vesicle distribution. These observations are consistent with previous results that a minimal difference in length between neurites is required for the extension of an axon (Dotti & Banker, 1987; Goslin & Banker, 1989; Yamamoto *et al*, 2012). The recruitment of constitutively active kinesins to PPVs before neuronal polarity is established was sufficient to induce the extension of supernumerary axons. This result suggests that increasing the supply of PPVs also promotes their exocytosis, which is required for axon extension to occur, implying that vesicle transport and exocytosis are coupled (Jacobson *et al*, 2006; Pfenninger, 2009; Jenkins *et al*, 2012; Deng *et al*, 2014).

To identify a link between PPV transport and the downstream exocytic machinery, we examined the role of Rab10, which regulates both, transport and exocytosis (Wang *et al*, 2011; Deng *et al*, 2014). The interaction of Rab10 with the scaffolding protein Jip1 links PPVs to Kinesin-1 and mediates their anterograde transport (Deng *et al*, 2014). Activation of Rab10 on PPVs by recruiting the catalytic domain of the Rab10 GEF DENND2B was sufficient to induce the extension of supernumerary axons, whereas activation at the plasma membrane did not have this effect. This result suggests that Rab10 not only mediates the transport of PPVs but is also involved in later steps leading to membrane expansion, consistent with previous results (Wang *et al*, 2011; Deng *et al*, 2014). Our results indicate that Rab10 is linked to PPV exocytosis by Arhgef7 and Cdc42b. Both were shown before to promote exocytosis in differentiating neurons upstream of the exocyst subunit Exo70 (Dupraz *et al*, 2009; López Tobón *et al*, 2018; Ravindran & Püschel, 2023). Their biochemical and functional interaction demonstrates that Rab10 acts upstream of Arhgef7 to promote the extension of axons. Rab10 binds to Arhgef7 and activates it in a heterologous system. Expressing a combination of constitutively active mutants and knockdown constructs in neurons confirmed that Rab10 acts upstream of Arhgef7, Cdc42b and Exo70 to induce the extension of supernumerary axons.

If the interaction of Rab10 and Arhgef7 stimulates PPV exocytosis, activating Arhgef7 specifically on PPVs should be sufficient to promote axon extension. Indeed, Arhgef7 activity was required on PPVs but not at the plasma membrane, similar to Rab10. The Cdc42 GEF Fgd1 had the opposite preference and induced supernumerary axons only when recruited to the plasma membrane but not when localized to PPVs. Thus, the function of the two Cdc42 GEFs requires a distinct subcellular localization to promote extension (Garvalov *et al*, 2007; Woo *et al*, 2019; Ravindran & Püschel, 2023). Since Cdc42 is not detectable on isolated PPVs (Deng *et al*, 2014), it is likely that it is activated in the growth cone by the delivery of vesicles. Together, these results support a model in which Rab10 activates Arhgef7 on PPVs, which in turn stimulates Cdc42b to engage Exo70 and promote PPV exocytosis in the growing neurite (Fig 8).

The essential role of exocyst-dependent exocytosis was confirmed also for the induction of supernumerary axons after actin depolymerization by LatA. The regulation of F-actin dynamics determines the rate of microtubule growth that drives axon extension (Witte *et al*, 2008; Dupraz *et al*, 2019; Burute *et al*, 2022; Schelski & Bradke, 2022; Lin *et al*, 2026). The depolymerization of F-actin removes the restriction on microtubule growth and induces the formation of supernumerary axons (Bradke & Dotti, 1999; Dupraz *et al*, 2019; Lin *et al*, 2026). The knockdown of Exo70 blocked this effect. Microtubules not only provide the physical backbone for axon elongation but also enable the anterograde transport of PPVs by Kinesin-1 (Deng *et al*, 2014). The supply of PPVs for membrane expansion then allows the further growth of microtubules (Pfenninger *et al*, 2003; Jacobson *et al*, 2006; Witte *et al*, 2008; Schelski & Bradke, 2017). The movement of Kinesin-1 along microtubules can modulate their dynamics and could further enhance anterograde transport (Andreu-Carbó *et al*, 2022). The requirement of Exo70 for the induction of supernumerary axons by F-actin depolymerization confirms that cytoskeletal dynamics and membrane expansion are inter-dependent processes that are both required for axon extension (Schelski & Bradke, 2017).

The biased distribution of PPVs to the longest neurite and the induction of supernumerary axons by an increased supply of PPVs raise the question how a difference in neurite length and the polarization of transport are linked. Different signaling mechanisms have been suggested for the neurite length-dependent regulation of neuronal polarity.

Anterograde transport and retrograde diffusion can account for the length-dependent accumulation of Shootin1 that interacts indirectly with actin filaments and promotes axon extension (Toriyama *et al*, 2010). Another model is based on long-range Ca^2+^ waves that activate RhoA in the soma, resulting in an inhibition of minor neurites by the diffusion of Rho kinase (Takano *et al*, 2017). Both models address a length-dependent regulation of actin-linked processes but not how vesicle trafficking becomes polarized. A frequency-encoded signal depending on the molecular motors that mediate bidirectional transport was described for detecting neurite length in sensory neurons (Rishal *et al*, 2012). This model indicates an important role for intracellular transport itself in sensing neurite length.

Simulations based on a model for vesicle transport in polarizing neurons suggested that the increased exocytosis of vesicles that sustains neurite growth can itself result in polarized vesicle trafficking to the longest neurites (Humpert *et al*, 2021). In this model, vesicles are preferentially transported into the longest neurite once a sufficient difference in neurite length is established. The mathematical framework introduced here to describe a coupling between neurite length and vesicle transport identifies a critical length asymmetry associated with a degenerate pitchfork bifurcation (Fig 2G), providing a minimal dynamical description of the onset of a polarization in vesicle transport. We also explored an extended formulation, in which neurite length asymmetry evolves dynamically in response to the polarization of vesicle transport. This extension provides a framework for investigating the reciprocal coupling between vesicle transport and neurite growth. The extended model suggests that the symmetry-breaking mechanism identified in the minimal model may be part of a larger feedback loop governing neuronal polarization, consistent with models in which neurite growth, polarity signaling, and cytoskeletal dynamics mutually reinforce one another (Schelski & Bradke, 2017). A more detailed analysis of this framework in the future may help to elucidate how the interaction of these processes translates into the establishment of a polarized neuronal morphology. The inclusion of a link between vesicle transport and neurite growth implies a coupling of vesicle abundance and fusion rate, i.e. a mechanism by which an increased vesicle supply facilitates exocytosis. The regulation of Arhgef7 by Rab10 on PPVs could mediate this link between vesicle supply and exocytosis, and, thereby, the polarization of vesicle transport to the longest neurite. However, it is difficult to directly show that Rab10 and Arhgef7 mediate the polarization of vesicle transport since Rab10 is also required for transport and both, Rab10 and Arhgef7, are essential for axon extension (Wang *et al*, 2011; Deng *et al*, 2014; López Tobón *et al*, 2018; Ravindran & Püschel, 2023).

Taken together, experimental results and theoretical considerations support a mechanism in which an initial neurite length asymmetry leads to a bias in vesicle transport toward the longer neurite, thereby establishing a positive feedback loop. We identify an interaction between Rab10 and Arhgef7 that promotes Cdc42b activation and is essential for axon extension, which could link vesicle transport and exocytosis.

## Materials and Methods

### Experimental model and subject details

All procedures involving animals complied with the regulations of the North Rhine-Westphalia State Environment Agency (Landesamt für Natur, Umwelt und Verbraucherschutz). Rats were housed in the animal facility of the Institute for Integrative Cell Biology and Physiology at the University of Münster under standardized conditions, including a 12-h light/dark cycle (07:00–19:00), an ambient temperature of 23°C, and unrestricted access to food and water. Timed mating was performed in-house. On embryonic day 18 (E18), pregnant rats were euthanized by decapitation after isoflurane anesthesia and embryos collected for the preparation of primary neuronal cultures. Neurons obtained from all embryos within a litter were combined, without distinguishing between male and female embryos.

### Neuronal cell culture and transfection

Neuronal culture and transfection were performed as previously described (Ravindran & Püschel, 2023). Briefly, primary cortical or hippocampal neurons were obtained from E18 rat embryos and transfected using calcium phosphate–mediated co-precipitation.

Following dissection, cortices or hippocampi were transferred to HBSS (Invitrogen) and enzymatically dissociated with 0.25% trypsin–EDTA (Invitrogen) for 8 min. The tissue was gently triturated in DMEM containing 2 mM glutamine, 50 U/ml gentamicin, and 10% FCS and filtered through a 40-µm cell strainer (Sarstedt). 13,000 cells/cm² were plated on glass coverslips or culture dishes (Knittel GLASS, Ibidi) coated with poly-L-ornithine (10 µg/ml; Sigma-Aldrich). 52,000 cells/cm² were plated for cultures with microstructures. After 2 h in DMEM, the medium was replaced with Neurobasal Medium (NBM) supplemented with 2 mM glutamine, 50 U/ml gentamicin, and 2% B27 (Invitrogen). For transfection, the culture medium was replaced by Opti-MEM 3.5 h after plating, followed by addition of the DNA– calcium phosphate mixture. Cells were incubated for 30 min at 37°C in 5% CO₂, the transfection mixture was removed, neurons were washed for 10 min with 1 ml of Opti-MEM pre-equilibrated at 37°C and 10% CO₂ and NBM added. Unless otherwise indicated, cortical and hippocampal neurons were transfected at 0 d.i.v. and fixed on 3 d.i.v..

### Plasmids

psfGFP-C1 (plasmid #54579; Addgene) was obtained from Geoffrey Waldo. psfGFP-C2, psfGFP-Arhgef7 and pcDNA3-HA-Arhgef7 have been described before (López Tobón *et al*, 2018; Ravindran & Püschel, 2023), pcDNA3-HA-Arhgef7-W200K,Y599E, sfGFP-Arhgef7-W200K, sfGFP-Arhgef7-Y599E, sfGFP-Arhgef7-W200K,Y599E (Marston *et al*, 2020) were generated by site-directed mutagenesis using Phusion High-Fidelity DNA Polymerase (Thermo Fisher Scientific). Primers for the mutagenesis are listed in Table S1. pcDNA6.2-H2B-mRFP, pCDNA-6-2-GW, pcDNA6.2-H2B-mRFP-miR-Cdc42b, pcDNA6.2-H2B-mRFP-miR-Arhgef7, pcDNA6.2-H2B-mRFP-miR-Exo70 and eGFP-Cdc42b have been described before (Ravindran & Püschel, 2023; Di Meo *et al*, 2024). A miRNA directed against Rab10 was generated using the BLOCK-iT Pol II miRNA Expression Vector Kit (Table S1). sfGFP-Rab10-res was generated by site-directed mutagenesis using Phusion High-Fidelity DNA Polymerase (Thermo Fisher Scientific) and the primers listed in Table S1 to introduce two mutations in the binding sites for respective miRNA. sfGFP-Rab10, sfGFP-Rab10 Q68L, HA-Rab10, and HA-Rab10 S18V were generated using the primers listed in Table S1. eGFP-Cdc42b, eGFP-DmrC and myr-mCherry-2xDmrA have been described before (Ravindran & Püschel, 2023; Kundu & Püschel, 2026). 5’UTR-2xDmrA-mCherry-Rab10, 5’UTR-2xDmrA-mCherry-VAMP4, eGFP-Arhgef7_DHPH-DmrC, eGFP-Fgd1_DHPH-DmrC, eGFP-DENND(2B)-DmrC were generated from pHet1 (TakaraBio) as previously described (Woo *et al*, 2019; Kumar *et al*, 2022) The DENN domain of DENND2B was obtained from a cDNA clone in pBluescriptR (BC036655-TCH1003-GVO-TRI, BioCat). Kif5C_ac-eGFP-DmrC and Kif1A_ac-eGFP-DmrC were generated from pEYFP-KIF5C and Kif1A_ac-eGFP-DmrC, respectively, with primers listed in Table S1 to construct vectors for constitutively active Kinesins as previously described (Jacobson et al, 2006). pBa.Kif1a 1-396.GFP was a gift from Gary Banker and Marvin Bentley (Addgene plasmid # 45058), pEYFP-KIF5C was a gift from Anne Stephenson (Smith *et al*, 2006). pGEX-PAK-PBD has been described before (Bagrodia *et al*, 1998).

### Immunofluorescence staining

Immunofluorescence staining of cultured neurons was carried out as described before (Ravindran & Püschel, 2023). Neurons were fixed with 4% paraformaldehyde and 4% sucrose in 1xPBS for 15 min at 37°C and quenched in 50 mM ammonium chloride for 10 min at room temperature. Cells were permeabilized with 0.1% Triton X-100 in 1× PBS for 3 min and incubated for 1 h at room temperature in blocking buffer (2% normal goat serum, 2% bovine serum albumin, and 0.2% fish gelatin in PBS). Primary antibodies, diluted in 10% blocking buffer, were applied overnight at 4°C. Following four washes with 1× PBS, cells were incubated with the appropriate Alexa Fluor–conjugated secondary antibodies for 30 min at room temperature. Coverslips were mounted using Mowiol (Sigma-Aldrich).

### Transfection of HEK293T cells and Western blot

HEK 293T cells were transfected using calcium phosphate co-precipitation. Cells were collected in ice-cold PBS and lysed for 40 min at 4°C in buffer containing 50 mM Tris/HCl (pH 7.4), 150 mM NaCl, 1 mM DTT, 1.5 mM MgCl₂, 4 mM EDTA, 10% glycerol, 1% Triton X-100 and Complete protease inhibitor (Merck). Where appropriate, tyrosine and serine/threonine phosphatase inhibitors (Sigma-Aldrich) were included in the lysis buffer. Proteins were resolved by SDS–PAGE and transferred onto PVDF membranes.

Membranes were blocked for 1 h at room temperature in Tris-buffered saline and 0.1% Tween-20 (TBS-T) containing 5% non-fat dry milk. Primary antibodies, diluted in blocking buffer, were applied overnight at 4°C. After several washes with TBS-T, membranes were incubated with HRP-conjugated secondary antibodies for 1 h at room temperature.

Chemiluminescent signals were detected using an enhanced chemiluminescence system (Interchim) and captured with a ChemiDoc MP imaging system (Bio-Rad).

### Biochemistry

For pulldown experiments, GST-tagged proteins were expressed in Escherichia coli BL21 cells. GST-fusion proteins were coupled to Glutathione-sepharose 4B beads (VWR) for 1 h at 4°C. Beads were washed (Tris/HCl (pH 7.4), 20 mM, NaCl 25 mM, DTT 0.1 mM, Complete protease inhibitor) and incubated with lysates of transfected HEK293T cells for 2 or 3 h at 4°C. The beads were washed (50 mM Tris/HCl (pH 7.4), 150 mM NaCl, 1 mM DTT, 1.5 mM MgCl_2_, with or without (for GTPase activity assay) 5 mM EDTA, 10% glycerol, 0.1% Triton X-100, Complete protease inhibitor). Bound proteins were eluted using 2× Laemmli buffer and analyzed by Western blot.

### Preparation of patterned microstructure

Glass-bottom dishes (ibidi) were plasma-cleaned at maximum instrument intensity for 5– 10 min (instrument follows) to prepare the glass surface for silanization. Immediately after plasma treatment, 3-(trimethoxysilyl)propyl methacrylate (Sigma Aldrich) was applied undiluted to the center of each dish and allowed to react for 2 min. The silane solution was subsequently removed, the surface rinsed once with isopropanol and allowed to dry. The hydrogel precursor was prepared using poly(ethylene glycol) diacrylate (PEGDA; molecular weight 700 Da, Sigma Aldrich) containing 1% (w/w) 2-Benzyl-2-(dimethylamino)-4′-morpholinobutyrophenone (DBMP, Sigma Aldrich) as the photoinitiator. The components were mixed thoroughly on a roller for 24 h to obtain a homogeneous precursor solution.

Microstructured hydrogel patterns were fabricated using a custom-built two-photon polymerization (2PP) setup, as described previously (Dzikonski *et al*, 2025). A small volume of the hydrogel precursor was added to each silanized dish and a coverslip was placed on top to confine the precursor and minimize fluid movement during patterning. Pattern geometries were generated in MATLAB and laser patterning was performed automatically in an array configuration. Following polymerization, the coverslip was removed and the patterned structures were washed three times with ethanol to remove residual precursor. The structures were stored in ethanol until use. Before cell seeding, the structures were washed three times with sterile PBS, coated with poly-L-ornithine in PBS as described above and then rinsed once with PBS. Culture medium was added and cells were seeded as described above.

### Numerical simulations

The dynamical system was solved numerically using the *solve_ivp* function from the SciPy library in Python, employing the Runge–Kutta method of order 5(4) (RK45). This adaptive-step integration method uses a fifth-order formula to advance the solution and an embedded fourth-order formula to estimate the local truncation error and adjust the step size accordingly. The system was integrated over the time interval *t* ∈ [0, 84], and the solution was evaluated at 1000 equally spaced time points, with relative and absolute error tolerances set to 10^−7^and 10^−9^, respectively, and the maximum integration step size restricted to 0.1.

### Image acquisition and analysis

Images were acquired on a Zeiss LSM 800 confocal laser scanning microscope with 405, 488, 568 and 647 nm laser lines, and processed using the Zeiss ZEN 2.3 (blue edition) software (Carl Zeiss MicroImaging, Jena, Germany) and ImageJ (NIH). Neuronal morphology was analyzed using a Zeiss Plan-Achromat 20x/0.4 objective. Brightfield images were captured with an iDS UI-1240ML-NIR-GL CMOS camera (IDS, Obersulm, Germany).

### Quantification

Neuronal polarity was assessed at 3 d.i.v. as described previously (Dhumale *et al*, 2018; Ravindran & Püschel, 2023). Neurons without acetylated tubulin–positive neurites were counted as unpolarized neurons without an axon, neurons with a single acetylated tubulin– positive neurite as polarized neurons with a single axon, and neurons with several acetylated tubulin–positive neurites as neurons with multiple axons (Witte *et al*, 2008). At least 100 neurons were evaluated per condition in each experiment, unless stated otherwise. The polarization index was calculated as the fluorescence intensity of the longest neurite (normalized to the length) divided by the intensity of the second longest neurite (normalized to the length) as described before (Grassi *et al*, 2015).

### Statistical analysis

Data are presented as means ± SEM, with each experiment conducted independently at least three times. The exact number of replicates is indicated in the figure legends. Unless stated otherwise, differences between two groups were evaluated using an unpaired *t*-test or a ratio paired *t*-test, when ratios were analyzed, whereas datasets involving more than two groups were analyzed by two-way ANOVA followed by Tukey’s multiple-comparisons test (GraphPad Prism). No statistical methods were used to determine the sample size.

The data were assumed to be normally distributed, although this assumption was not tested formally. Blinding was not performed. For all analyses performed, significance was defined as ns, P > 0.05; *P ≤ 0.05; **P ≤ 0.01; ***P ≤ 0.001; and ****P ≤ 0.0001. When not indicated, the conditions did not show a significant difference.

## Data Availability

Data sets generated from the current study are available from the corresponding author upon reasonable request.

## Acknowledgements

We thank Maria Wenning and Verena Stegemann for technical assistance. This work was supported by the Deutsche Forschungsgemeinschaft (DFG) through a grant to A.W.P. (PU 102/17-1).

## Author contributions

F.O.: conceptualization, data curation, formal analysis, investigation, visualization, methodology, and writing - original draft, review, & editing.

E.B.: data curation, formal analysis, investigation, visualization, methodology, software, and writing - original draft review & editing.

P.R.: data curation, formal analysis, investigation, methodology, writing - review & editing. T.P.: data curation, formal analysis, investigation, writing - review & editing.

A.W.P.: conceptualization, resources, data curation, supervision, funding acquisition, validation, visualization, project administration, writing - review and editing.

## Supplementary figures legends

**Figure S1. Exo70 is required for the induction of supernumerary axons by actin depolymerization (related to** Fig 1). Hippocampal neurons were transfected with control (Ctrl) or knockdown vectors for Exo70 expressing EmGFP (Figure 1E). 200 nM LatrunculinA (LatA) was added at 1 d.i.v. and axon extension analyzed at 3 d.i.v. with anti-acetylated tubulin (Cyan) and anti-MAP2 antibodies (magenta). Arrows mark axons. Scale bars: 20 µm.

**Figure S2. PPV distribution depends on neurite length (related to Fig 2).** (**A**) Length asymmetry (*x*(*t*), left) and vesicle transport polarization (*y*(*t*), right), are shown as a function of time. The length and the intensity of VAMP4 signals were determined for the longest and second longest neurite and length asymmetry and vesicle polarization were calculated according to 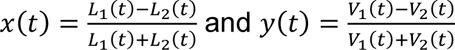 for each neuron (N=60 from n=3 biological replicates). (**B** - **E**) Solutions for the numerical integration of the coupled dynamical system, 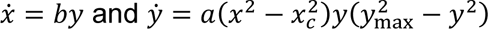, obtained by varying the transport parameter *a* (first and third rows) or the coupling coefficient *b* (second and fourth rows), while keeping the remaining parameters fixed. The left panels show the temporal evolution of normalized neurite length asymmetry, *x*(*t*), and the right panels show vesicle transport polarization, *y*(*t*). **(B)** Simulations with *x*_0_ = 0.01, *y*_0_ = 0.09, *x*_c_ = 0.15, and *y*_max_ = 1.0, with *b* = 0.20 for variation of *a* and *a* = 0.70 for variation of *b*. **(C)** Simulations with *x*_0_ = 0.01, *y*_0_ = 0.09, *x*_c_ = 0.30, and *y*_max_ = 0.80, using the same parameter settings for the respective parameter sweeps. **(D)** Simulations with *x*_0_ = 0.05, *y*_0_ = 0.04, *x*_c_ = 0.15, and *y*_max_ = 1.0, again varying *a* at fixed *b* = 0.20 and varying *b*at fixed *a* = 0.70. **(E)** Simulations with *x*_0_ = 0.05, *y*_0_ = 0.04, *x*_c_ = 0.30, and *y*_max_ = 0.80, using the same parameter settings for the respective parameter sweeps. Curves of different colors correspond to the parameter values indicated in the legends. The simulations illustrate how the critical length asymmetry, initial conditions, transport dynamics, and coupling strength influence the onset and temporal development of transport polarization and the associated growth of length asymmetry. Trajectories are shown over the simulated time interval; reaching transport saturation does not imply stabilization of length asymmetry.

**Figure S3. Increased PPV transport induces the extension of supernumerary axons (related to Fig 3).** (**A, B**) Cortical neurons were co-transfected with the indicated vectors (K1A/K5C_ca-eGFP-DmrC and DmrA-mCh-Rab10, Figure 3B and D). Solvent (DMSO, control) or heterodimerizer (0.25 nM) was added at 22 or 34 h of culture for 30 minutes. Neurons were analyzed at 3 d.i.v. by staining with anti-acetylated tubulin (cyan) and anti-MAP2 antibodies (magenta). Arrows mark axons. Scale bars: 20.

**Figure S4. Rab10 activation on PPVs promotes axon formation (related to** Fig 4). Cortical neurons were co-transfected with the indicated vectors (DmrC or DENND(2B)-DmrC, DmrA-Rab10, DmrA-VAMP4 or myr-DmrA, Figure 4B). Solvent (DMSO, control) or heterodimerizer was added at 22 h of culture for 30 minutes. Neurons were analyzed at 3 d.i.v. by staining with anti-acetylated tubulin (cyan) and anti-MAP2 antibodies (magenta). Arrows mark axons. Scale bars: 20 µm.

**Figure S5. Arhgef7 acts upstream of Cdc42b and Exo70 to promote axon extension (related to Fig 5).** (**A**) Hippocampal neurons were transfected with vectors for Arhgef7, Arhgef7-W200K, Arhgef7-Y599E or Arhgef7-W200K,Y599E and GFP (top panels, Figure 5) and analyzed at 3 d.i.v. by staining with anti-acetylated tubulin (cyan) and anti-MAP2 antibodies (magenta). Scale bar: 20 µm. (**B**) The percentage of neurons without an axon (0), with a single axon (1) and with multiple axons (>1) was quantified for (A) (n = 3 independent experiments that are biological replicates, N > 300 neurons). (**C, D**) Hippocampal neurons were co-transfected with control (Ctrl) or RNAi vectors directed against Cdc42b expressing H2B-RFP (right panels) and vectors for sfGFP, sfGFP-Arhgef7 or sfGFP-Arhgef7-ca (C, Figure 5D), or with control (Ctrl) or RNAi vectors for Exo70 expressing H2B-RFP, and vectors for sfGFP, sfGFP-Arhgef7 or sfGFP-Arhgef7-ca (D, Figure 5F). Neurons were analyzed at 3 d.i.v. by staining with anti-acetylated tubulin (cyan) and anti-MAP2 antibodies (magenta). Arrows mark axons. Scale bars: 20 µm. Values are means ± SEM. **(B)**: two-way ANOVA.

**Figure S6. Rab10 interacts with and activates Arhgef7 to promote PPV exocytosis (related to Fig 6).** (**A**) HEK 293T cells were transfected with vectors for eGFP-Rab10 or RNAi-resistant eGFP-Rab10 (Rab10-res), and control (emGFP) or RNAi vectors for Rab10 expressing in emGFP (emGFP-RNAi). The efficiency of the knockdown was analyzed by Western blot (WB) using an anti-GFP antibody. (**B**) The expression of Rab10, normalized to the expression of emGFP, was quantified for (A) (n = 3 independent experiments that are biological replicates). (**C**) Hippocampal neurons were co-transfected with control or RNAi vectors for Rab10 expressing in emGFP (right panels, black), and vectors expressing sfGFP or RNAi-resistant sfGFP-Rab10 (Rab10-res). Neurons were analyzed at 3 d.i.v. with anti-acetylated tubulin (cyan) and anti-MAP2 antibodies (magenta). (**D**) The percentage of neurons without an axon (0), with a single axon (1) and with multiple axons (>1) was quantified for (C) (n = 4 independent experiments that are biological replicates, N > 400 neurons). (**E**) Hippocampal neurons were co-transfected with control (Ctrl) or RNAi vectors for Cdc42b expressing H2B-RFP, and vectors for sfGFP, sfGFP-Arhgef7 or sfGFP-Arhgef7-ca (see Figure 6D). Axon extension was analyzed at 3 d.i.v. with anti-acetylated tubulin and anti-MAP2 antibodies. (**F**) Hippocampal neurons were co-transfected with control (Ctrl) or RNAi vectors for Exo70 expressing H2B-RFP, and vectors for sfGFP, sfGFP-Arhgef7 or sfGFP-Arhgef7-ca (Figure 6F). Neurons were analyzed at 3 d.i.v. by staining with anti-acetylated tubulin (cyan) and anti-MAP2 antibodies (magenta). Arrows mark axons. Scale bars: 20 µm. Values are means ± SEM. **(B)**: ratio paired Student’s t-test, **(D):** two-way ANOVA.

**Figure S7. Recruitment of Arhgef7 to PPVs promotes axon extension (related to Fig 7).** Cortical neurons were co-transfected with the indicted vectors (DmrC, Arhgef7_DHPH-DmrC or Fgd1_DHPH-DmrC, Figure 7B, and DmrA-Rab10, DmrA-VAMP4 or myr-DmrA) as indicated. Vehicle (DMSO, control) or dimerizer was added for 30 min at 22 h of culture. Neurons were analyzed at 3 d.i.v. by staining with anti-acetylated tubulin (cyan) and anti-MAP2 antibodies (magenta). Arrows mark axons. Scale bars: 20 µm.

